## Supplementary figures and images for "Cell loss disrupts mechanical homeostasis to drive retinal pigment epithelium ageing-like phenotype *in vitro*"

### Sup. Fig. 1

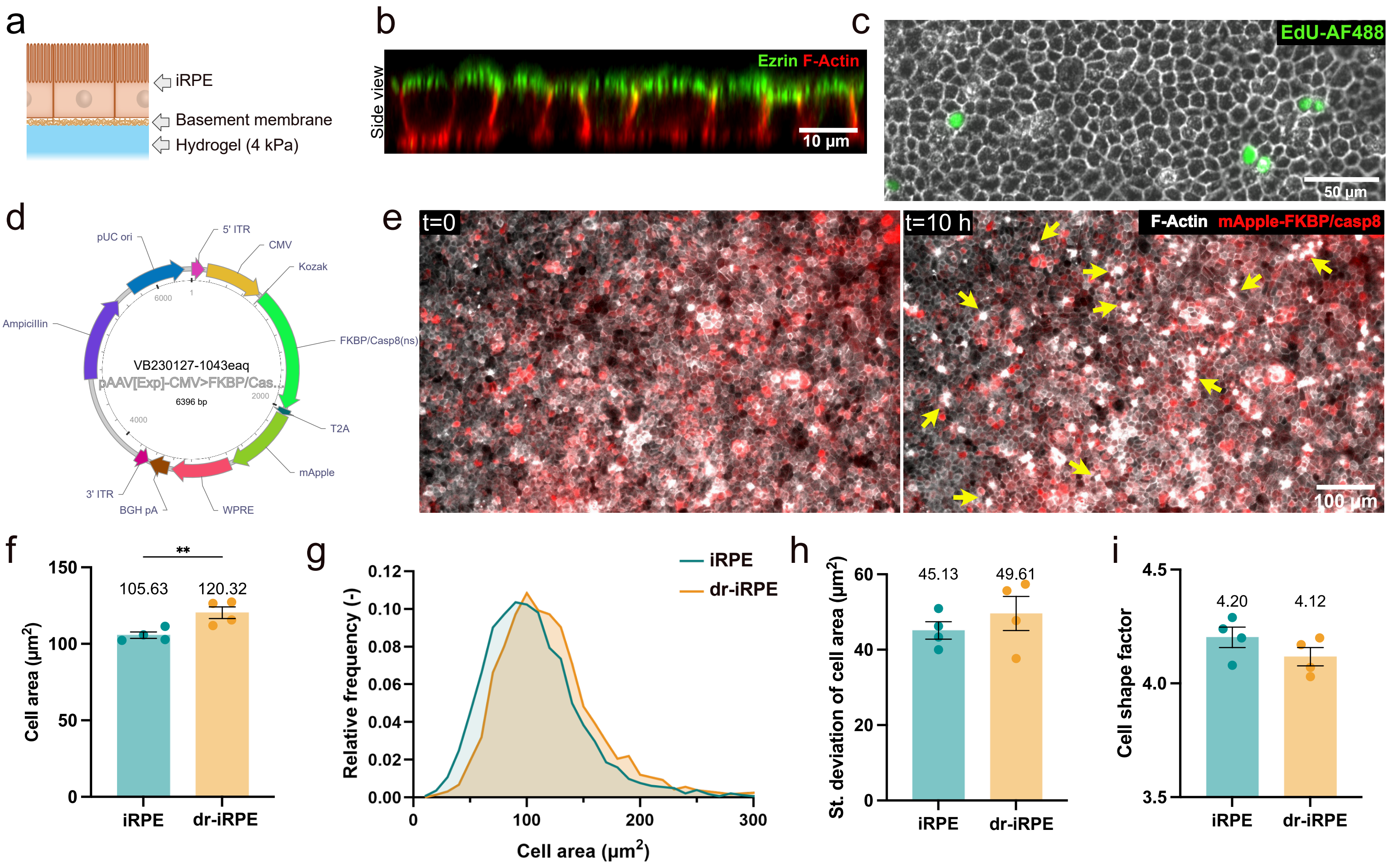

### Sup. Fig. 2

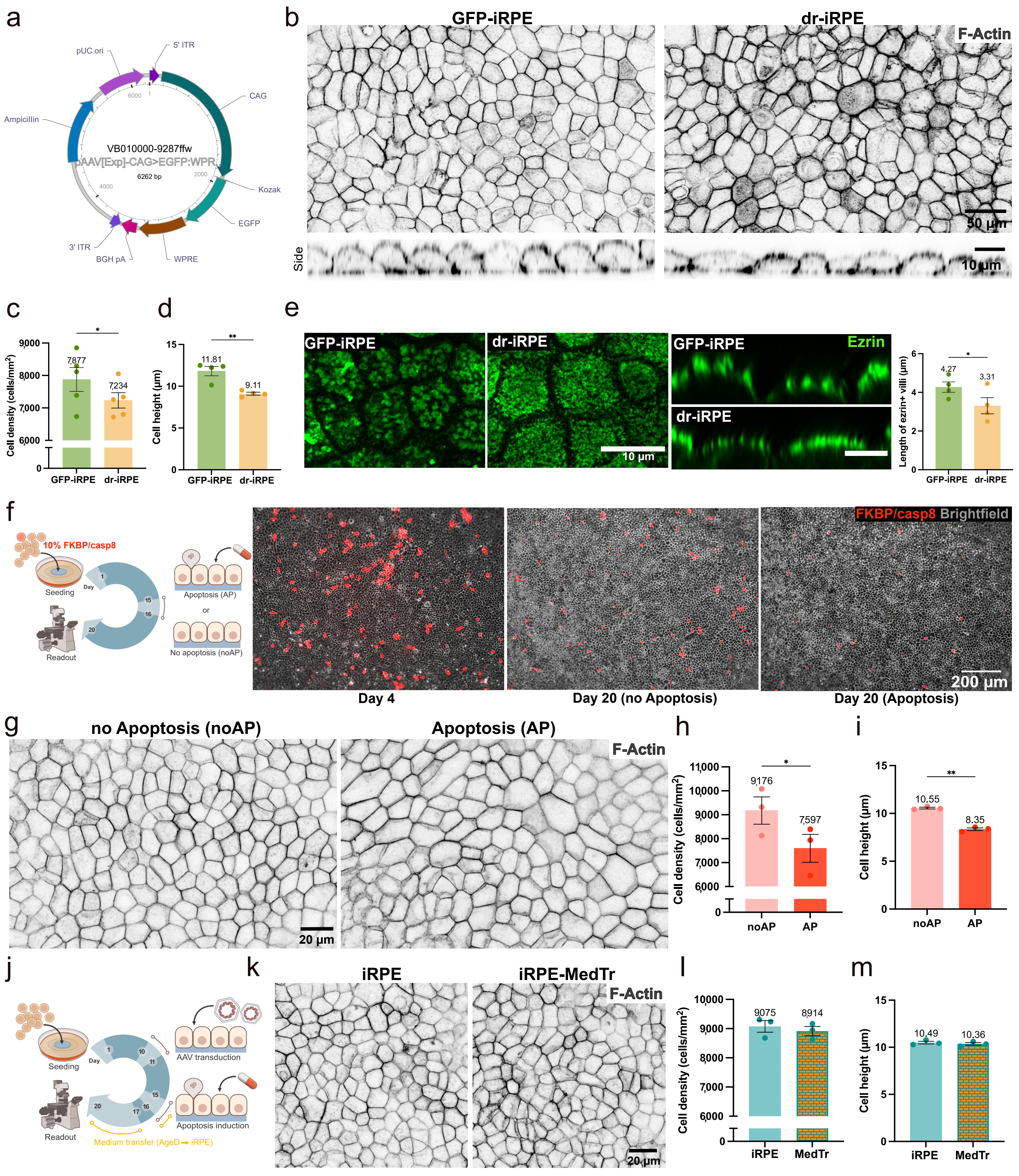

### Sup. Fig. 3

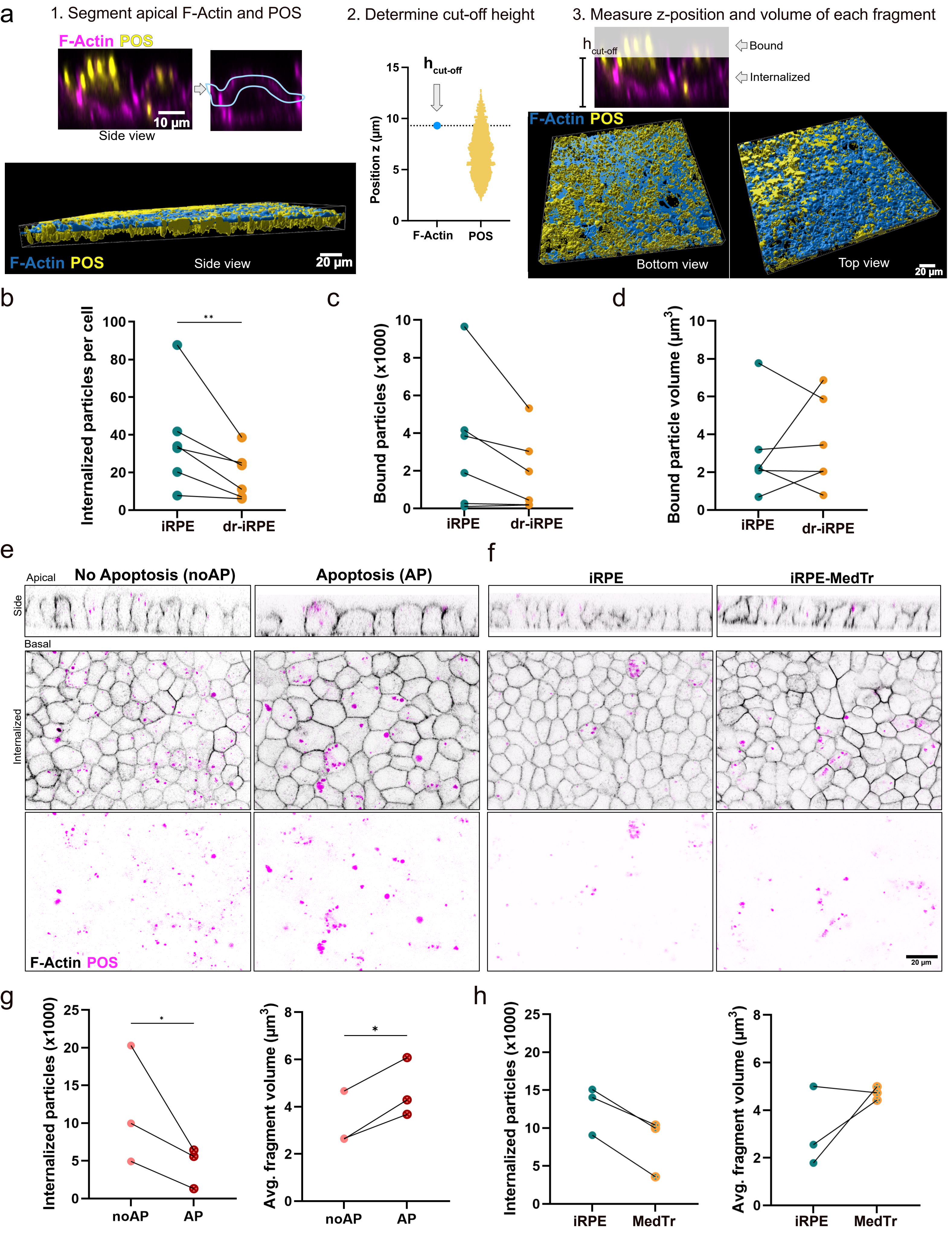

### Sup. Fig. 4

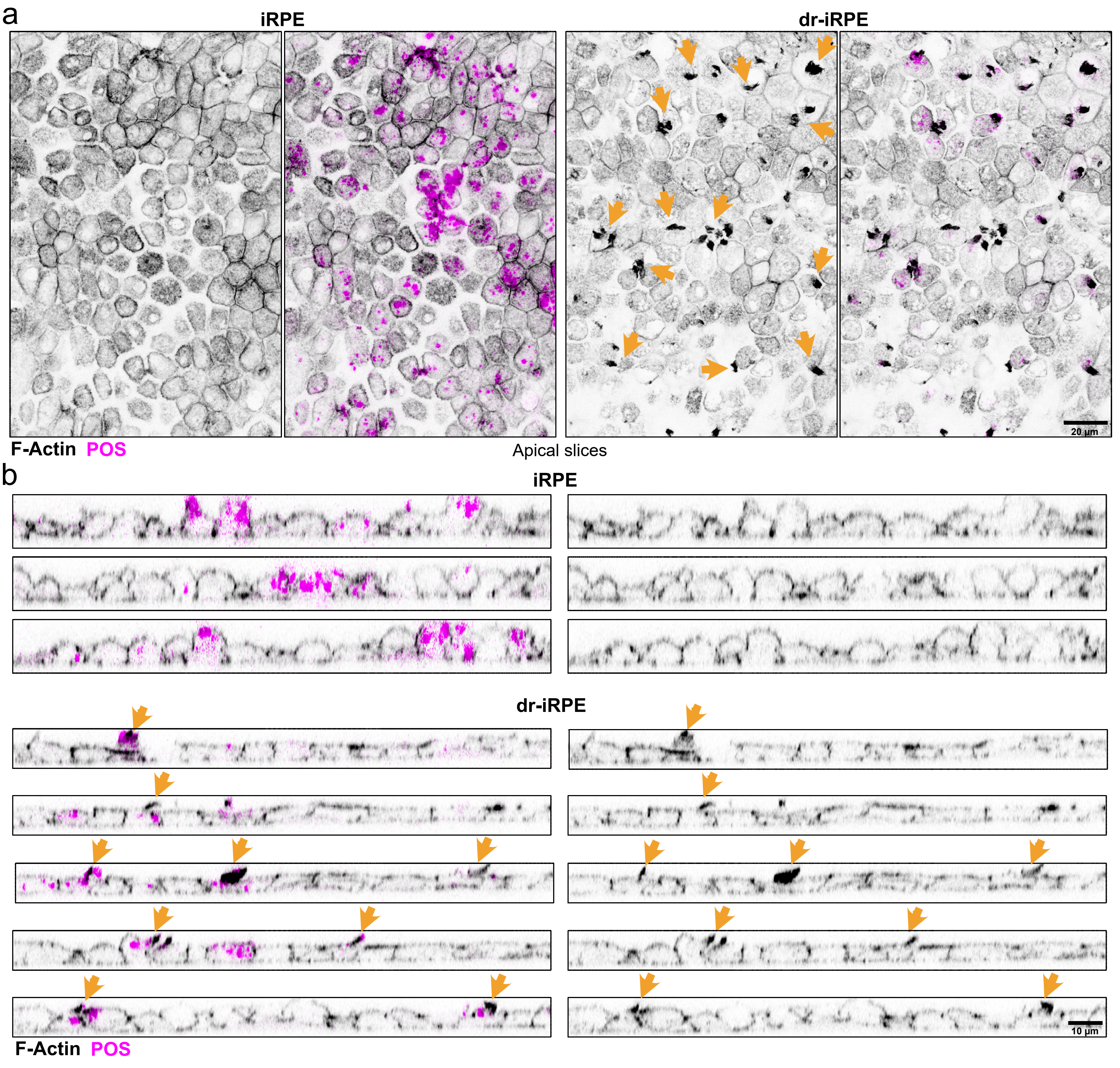

### Sup. Fig. 6

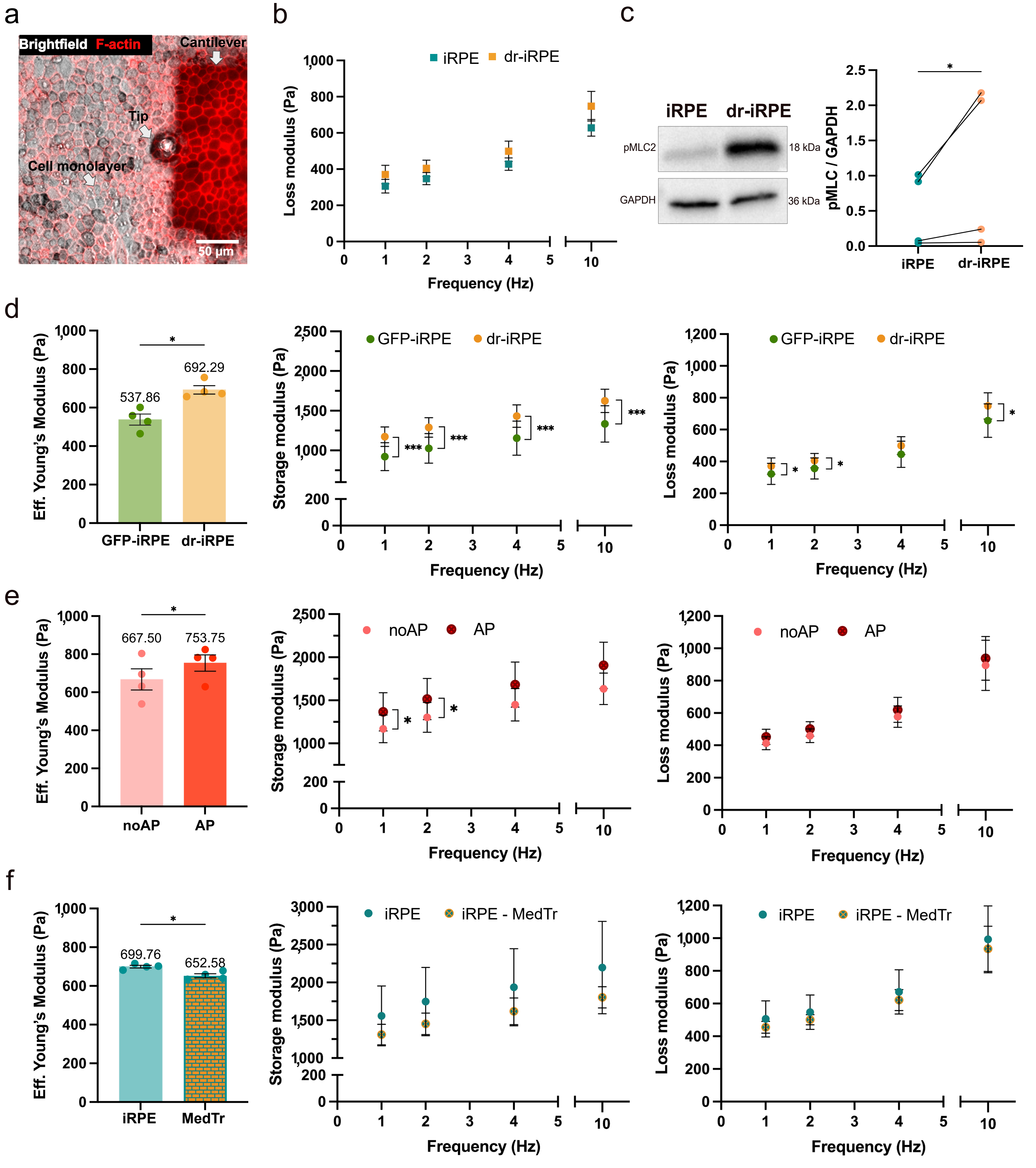

### Sup. Fig. 7

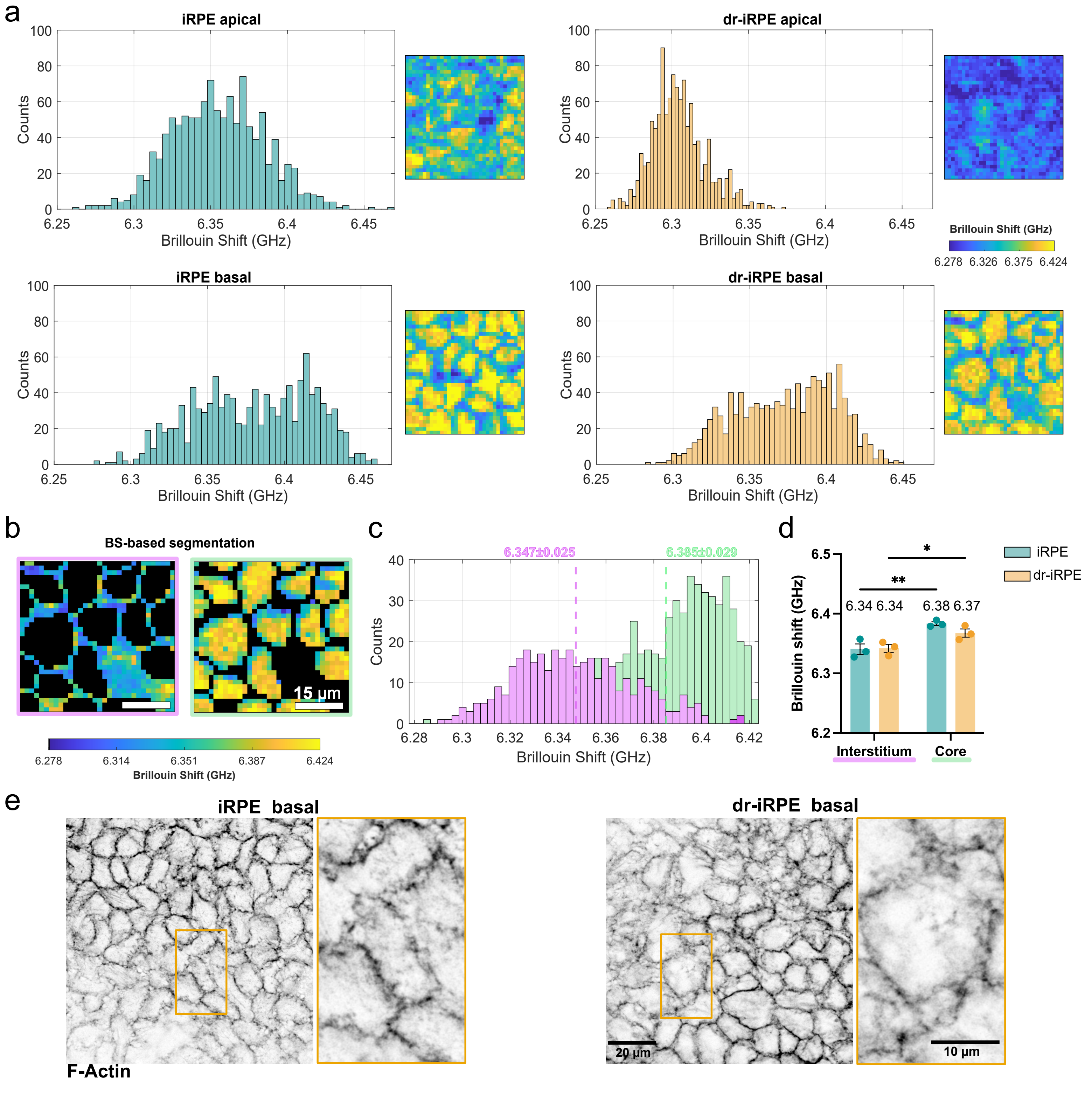

### Sup. Fig. 8

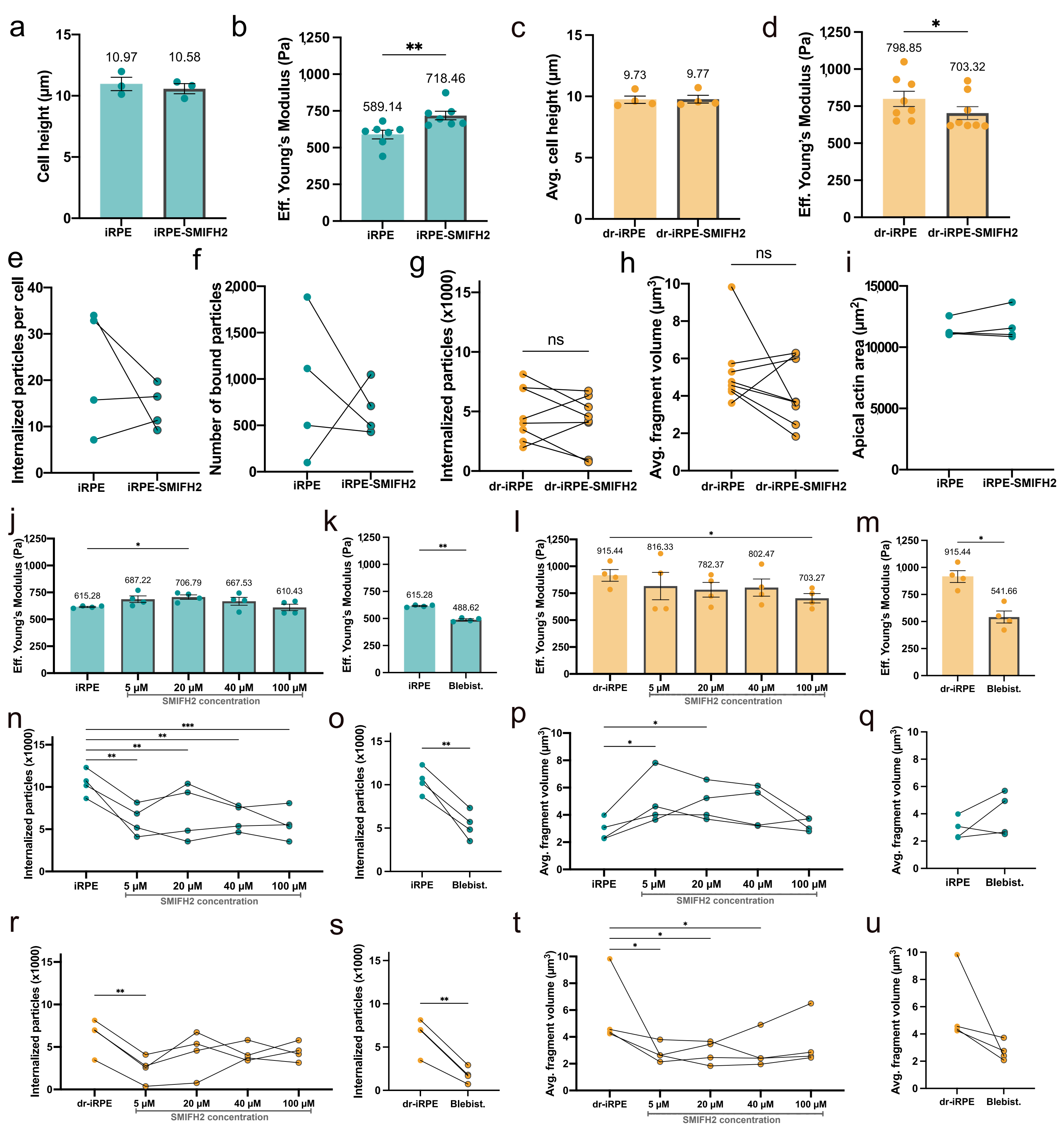

### Sup. Fig. 9

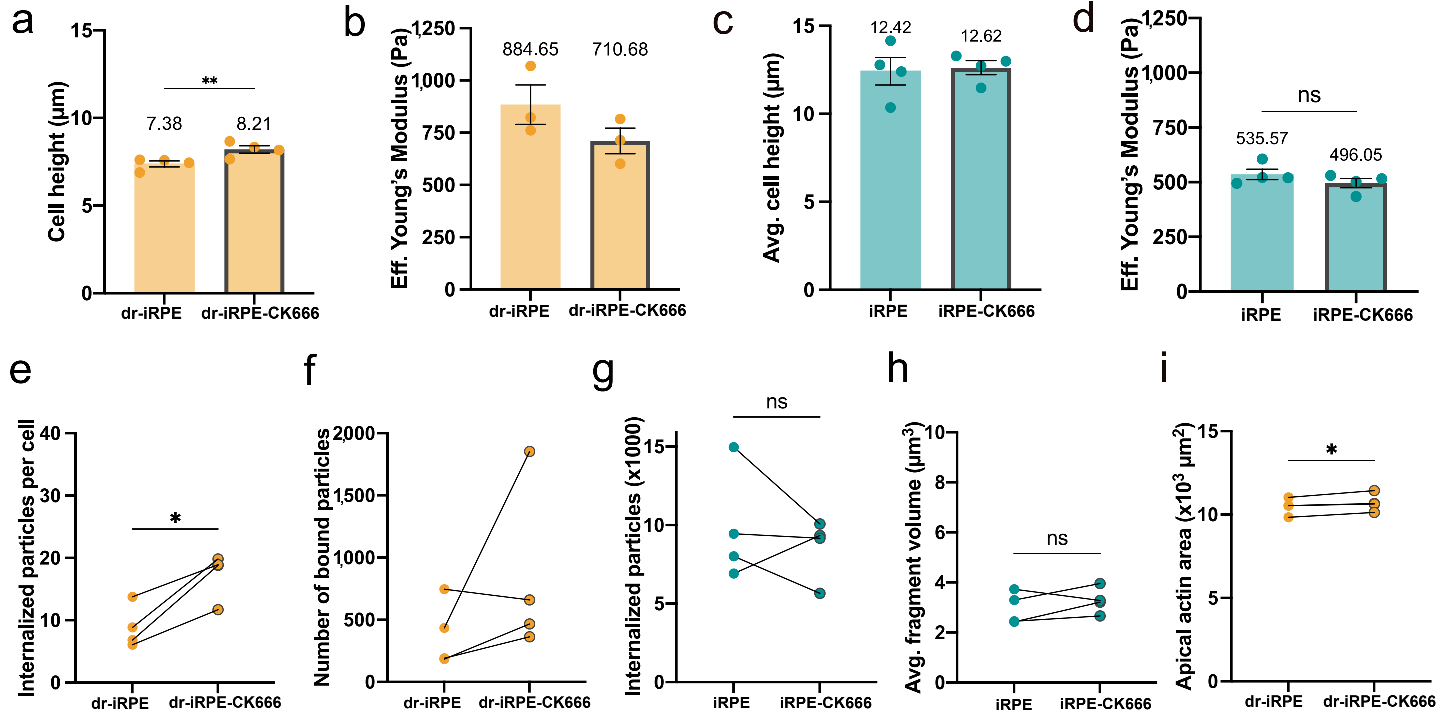
