## Supplementary material for "Cell loss disrupts mechanical homeostasis to drive retinal pigment epithelium ageing-like phenotype *in vitro*": Sup. Fig. 5

a

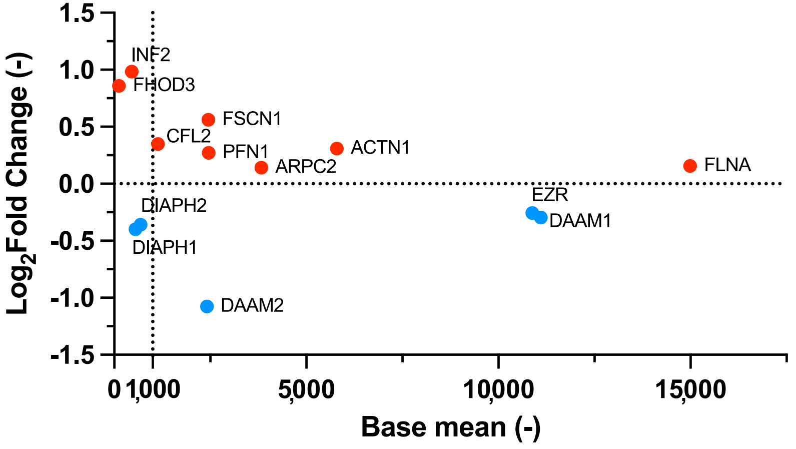


b

### Overlapping DEGs with reported age-related changes in humans

| Gene symbol | Protein name | Direction change |
|---|---|---|
| **Cai H. et al. 2012 (DOI:10.3389/fnagi.2012.00008)** | | |
| **DKK1** | dickkopf WNT signaling pathway inhibitor 1(DKK1) | up |
| **LAMA3** | laminin subunit alpha 3(LAMA3) | up |
| **TYMS** | thymidylate synthetase(TYMS) | up |
| **PLEKHG4** | pleckstrin homology and RhoGEF domain containing G4(PLEKHG4) | down |
| **SCN2B** | sodium voltage-gated channel beta subunit 2(SCN2B) | down |
| **Butler J. M. et al. 2021 (DOI: 10.1111/jcmm.16569)** | | |
| **AKAP6** | A-kinase anchoring protein 6(AKAP6) | up |
| **BMP8B** | bone morphogenetic protein 8b(BMP8B) | up |
| **CCDC51** | coiled-coil domain containing 51(CCDC51) | up |
| **CTSV** | cathepsin V(CTSV) | up |
| **DUSP6** | dual specificity phosphatase 6(DUSP6) | up |
| **KL** | klotho(KL) | up |
| **SLC6A20** | solute carrier family 6 member 20(SLC6A20) | up |
| **TNFRSF10C** | TNF receptor superfamily member 10c(TNFRSF10C) | up |
| **TRPV4** | transient receptor potential cation channel subfamily V member 4(TRPV4) | up |
| **CPAMD8** | C3 and PZP like alpha-2-macroglobulin domain containing 8(CPAMD8) | down |
| **CSRP2** | cysteine and glycine rich protein 2(CSRP2) | down |
| **DCT** | dopachrome tautomerase(DCT) | down |
| **GAS1** | growth arrest specific 1(GAS1) | down |
| **TMEM150C** | transmembrane protein 150C (TMEM150C) | down |

c

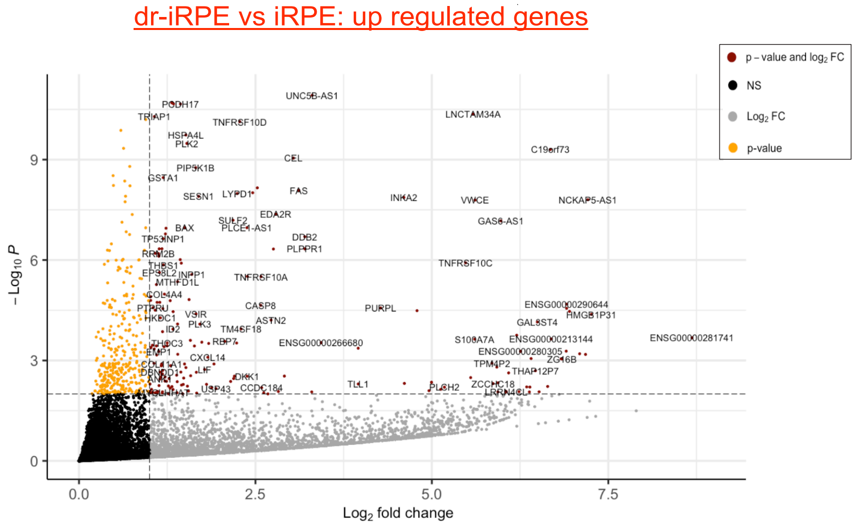


d

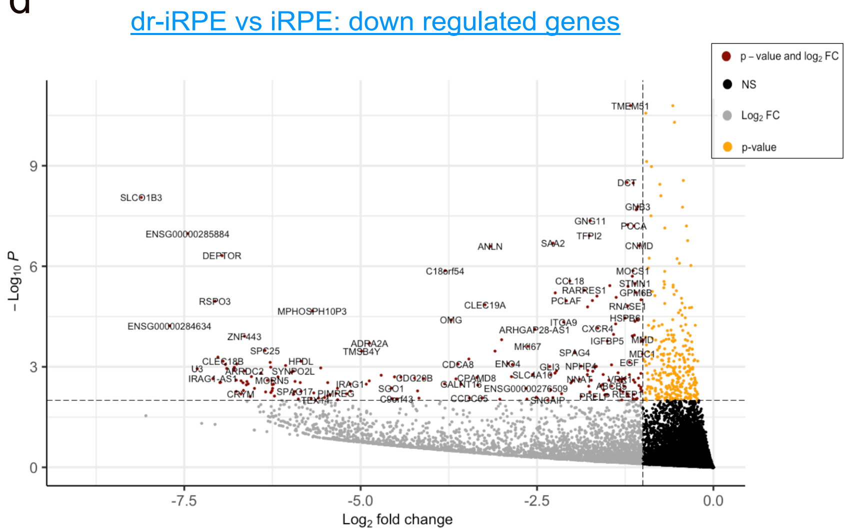
